# Generation of a Public Dataset for Etiological Subtyping with the Genomic Cluster Engine

**DOI:** 10.64898/2026.11.18.767536

**Authors:** Jędrzej Kubica, Mauricio Moldes, Ben Busby

**Author notes:** These authors contributed equally (Jędrzej Kubica, Mauricio Moldes).

## Abstract

**Motivation:** Genetic diseases are complex phenomena that are currently defined by their clinical symptoms, rather than by the combinations of variants that give rise to them. Incomplete penetrance and phenotypic variability indicate that the genomic background contributes to the effect of variants. Recent advancements in genomics have produced vast amounts of data and created new opportunities to study genetic diversity across human populations. Much of this variation is inherited as recombination-defined contiguous segments, called haploblocks. We previously introduced a novel methodology for genomic hash encoding, the Genomic Cluster Engine, which groups individuals into clusters using phased haploblock sequences. By encoding each haploblock sequence as a compact hash, the characterization of genomic backgrounds, the subtyping of etiologies and the inference of local ancestry become fast and scalable.

**Results:** We have optimized the Genomic Cluster Engine pipeline for large genomic datasets through parallelization and improved memory management. We applied our pipeline to 2,548 individuals across 26 populations from the 1000 Genomes Project and produced a dataset of haploblock-based genomic hashes for the resulting clusters. The genomic hashes and clusters can be used for investigating variant effects in the context of their haplotype backgrounds or for genotype-phenotype association studies. This comprehensive dataset is now publicly available, providing a valuable resource for both academic and industry researchers worldwide.

**Availability:** The haploblock boundaries, sequences, clusters and genomic hashes for the 1000 Genomes Project are available under the CC0 license at https://data.haploblocks.org and on the Registry of Open Data on AWS https://registry.opendata.aws.

**Key Messages:**

- We optimized the Genomic Cluster Engine for large-scale datasets.
- We applied our pipeline to 1000 Genomes and generated a dataset of phased haploblock sequences and genomic hashes.
- This dataset is now publicly available for academic and industry researchers worldwide.

## Introduction

Genomics research has improved our understanding of disease mechanisms by studying associations between genetic variation and clinical phenotypes [1, 2, 3]. However, many healthy individuals carry pathogenic variants without developing disease [4, 5, 6]. Genetic diseases have been consistently classified by their symptoms, rather than the combinations of specific variants that give rise to them, which have made genotype-phenotype interactions difficult to resolve. Previous studies suggest that this phenotypic variability is determined not only by pathogenic variants, but also by complex interactions between variants [7, 8] and their broad genomic backgrounds [9, 10, 11, 12]. Although many computational methods have been developed to predict the functional impact of single variants, studying interactions between variants and genomic background remains difficult, as such analyses require large-scale cohorts and scalable algorithms [13, 14, 15]. Investigating the effect of a variant in the context of its genomic background using whole-genome sequences is infeasible at scale due to the computational complexity of pairwise sequence comparison.

In addition, integrating large-scale datasets is limited by restrictions on data sharing and privacy [16].

Human genetic diversity is shaped in large part by recombination, a process whereby DNA fragments are exchanged between homologous chromosomes [17, 18]. These fragments are exchanged at non-uniform rates across the genome: regions with higher recombination rates compared to their surroundings are called recombination hotspots [19, 20]. Between them lie haploblocks, regions characterized by highly conserved allele inheritance patterns [21, 22]. Studying different combinations of alleles within and across haploblocks provides a new opportunity to identify interactions between variants and genomic backgrounds that shape individual phenotypes.

We previously introduced the Genomic Cluster Engine [23], a novel methodology that groups individuals by shared haploblock sequences and generates genomic hashes (Figure 1).

**Figure 1.**
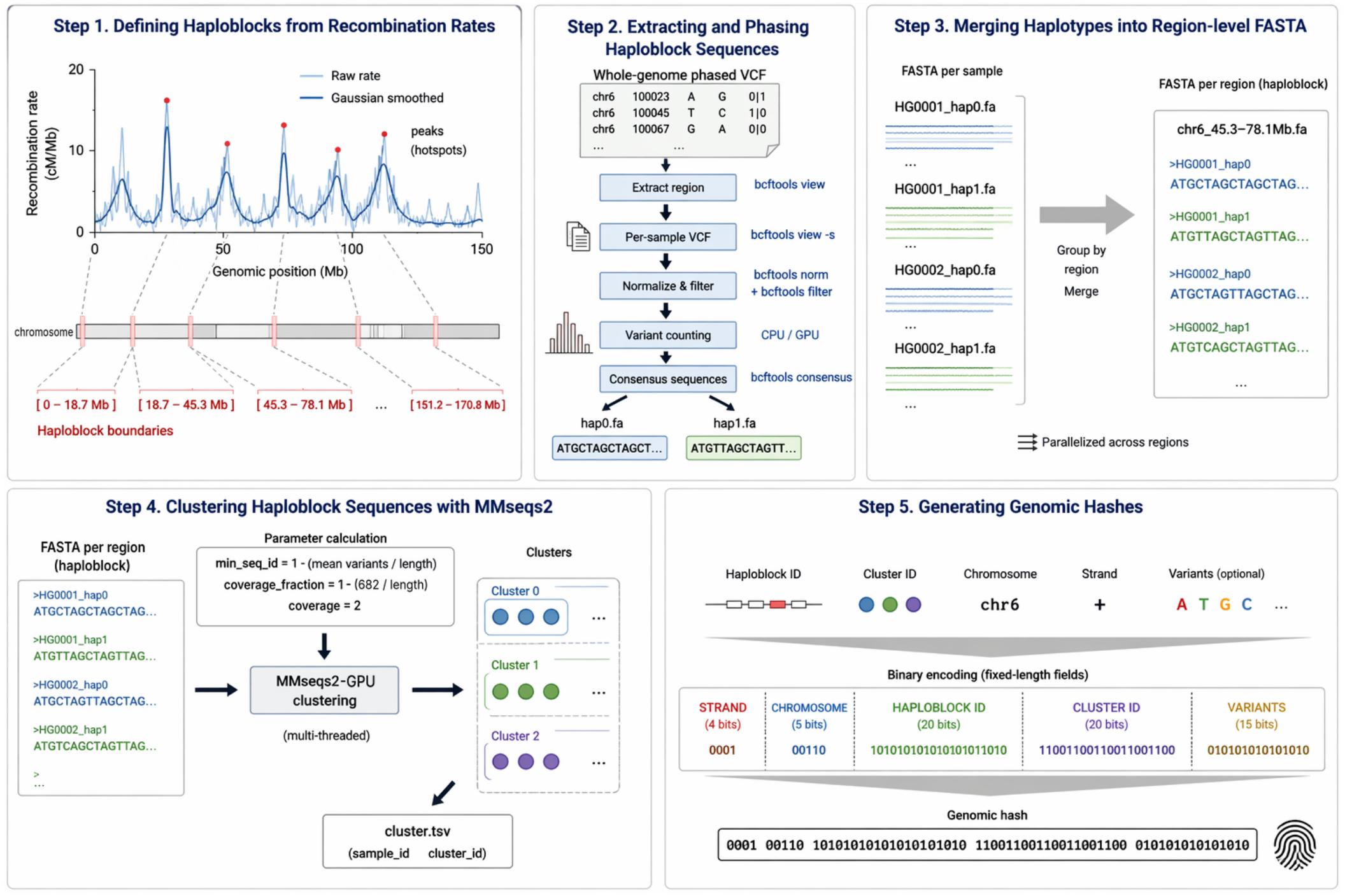
Schematic representation of the Genomic Cluster Engine. Step 1: Defining haploblocks using recombination hotspots (red) from Gaussian-smoothed recombination rates. Step 2: Extracting haploblock sequences from a phased VCF, divided by regions and by samples, and filtered based on minor allele frequency. Step 3: Merging phased haploblock sequences into one FASTA file per haploblock. Step 4: Clustering phased haploblock sequences using MMseqs2 with parameters *min*_*seq*_*id, coverage*_*fraction* and *coverage*. Step 5: Generating genomic hashes for phased haploblock sequences (with five fields: DNA strand, chromosome, haploblock, cluster, and optional variants of interest).

Rather than comparing whole genomes directly, the pipeline discretizes the genome into haploblocks and groups phased sequences within each haploblock. The pipeline comprises five steps: 1) defining haploblocks based on a recombination map, 2) extracting haploblock sequences from phased variant data (a phased VCF file), 3) merging phased haploblock sequences, 4) clustering phased haploblock sequences using MMseqs2 [24] with parameters *min*_*seq*_*id, coverage*_*fraction* and *coverage*, and 5) generating genomic hashes as binary strings (0s and 1s) that encode: DNA strand (4 characters), chromosome (10 characters), haploblock (20 characters ), and cluster (20 characters), with optional variants of interest (15 characters).

We used the Genomic Cluster Engine pipeline to generate a dataset with genomic hashes for the 1000 Genomes Project [25, 26]. We then analyzed the diversity of haplotypes within haploblocks and the resulting clusters in the population based on the genomic hashes.

## Methods

### Data

We used the empirically defined genome-wide recombination map with a 682-base-pair resolution from Halldorsson et al. [17] to approximate haploblock boundaries across the genome.

We obtained phased variant data for 2,548 individuals from 26 populations in the 1000 Genomes Project Phase 3 dataset (https://ftp.1000genomes.ebi.ac.uk) [25, 26]. The variant data has been mapped to the human reference genome sequence (GRCh38; https://www.ncbi.nlm.nih.gov), and phased with SHAPEIT2 [27].

### Pipeline optimization

We optimized the Genomic Cluster Engine pipeline, developed during the ELIXIR BioHackathon Europe 2025 and described in Kubica et al. [23]. The pipeline was dominated by I/O, data parsing, and sequential preprocessing, which limited GPU utilization. To address this, we included GPU integration across the pipeline, minimized CPU-GPU data transfer, increased computational density, and restructured earlier steps around GPU-native data representations. Clustering can now performed with the GPU implementation of MMseqs2 [28]. The pipeline is deterministic across CPU and GPU, including multithreading.

For high-performance computing infrastructures, we provide Docker and Singularity setups, and a configuration YAML file specifying inputs (recombination map, phased VCF, reference genome sequence), output locations for each step, and parameters (thread count, GPU usage, target chromosome). Each step of the pipeline can be run individually or via an orchestrator script that chains all steps.

### Pipeline execution

We executed the Genomic Cluster Engine pipeline using the recombination map and the phased variant data. The pipeline used the recombination map to identify haploblock boundaries, which were then used to divide the reference genome sequence into haploblock sequences. For each individual, common variants (minor allele frequency, *MAF >* 0.05) from the phased variant data were mapped to the haploblock sequences. MMseqs2 algorithm *easy-cluster* was used to group individuals’ haploblock sequences into clusters based on sequence similarity. The clusters indicated the individuals sharing a similar sequence within the haploblock. MMseqs2 parameters were chosen as follows: “target sequence identity” was defined as 1 *™ haploblock*_*mean*_*number*_*of* _*variants*, “coverage fraction” was defined as 1 *−* (682*/haploblock*_*length*) (682 bp was the resolution of the recombination map), and “coverage” was set to 2 (i.e. “query coverage”). The pipeline generated genomic hashes for all individuals’ haploblock sequences except for those corresponding to telomeric regions, which could not be clustered.

We generated the dataset on an Linux-based system with 128 CPU cores, 480 GB of system memory and NVIDIA A100 GPU.

## Results

We examined the diversity of haploblock sequences across the genome based on the clusters and the genomic hashes.

### Genome-wide structure of haploblocks

The reference genome sequence was divided into 39,141 haploblocks across autosomal chromosomes and chromosome X. We first examined how these haploblocks were distributed across chromosomes (Figure 2).

**Figure 2.**
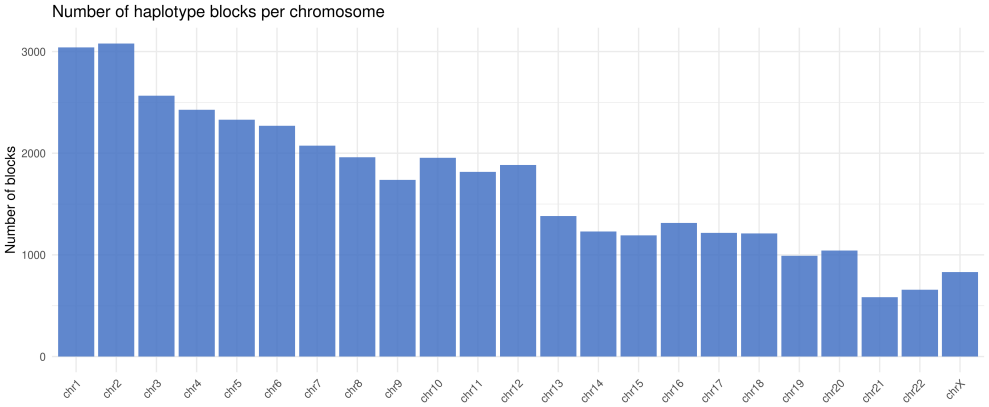
Haploblocks count per chromosome. The histogram represents the distribution of 39,141 haploblocks across 22 autosomes (chr1-22) and chromosome X (chrX), ordered by chromosome number.

Larger chromosomes tended to be discretized into more haploblocks: chromosomes 1 and 2 comprised over 3000 haploblocks, whereas chromosome 21 comprised just over 500 haploblocks. However, this relationship was not strict, for example chromosome 10 was divided into more haploblocks than the marginally longer chromosome 9 (chromosome 10: 133 Mb and 1968 haploblocks vs. chromosome 9: 138 Mb and 1748 haploblocks). Similarly, chromosome 16 was divided into more haploblocks than the longer chromosome 15 (chromosome 16: 90 Mb and 1322 haploblocks vs. chromosome 15: 102 Mb and 1210 haploblocks).

We then examined the distribution of haploblock lengths across the genome (Supp. Fig. 1). The median haploblock length was 48.4 kb, and most haploblocks (77.7%) fell between 10 kb and 100 kb. We also observed much shorter and longer haploblocks (outliers), ranging from 724 bp to 6.4 Mb.

We also examined the distribution of haploblock lengths by chromosome (Figure 3). Lengths varied moderately across autosomes, with median lengths between 29.7 (chromosome 22) and 58.8 kb (chromosome 4). In contrast, the median length of haploblocks in chromosome X (113.8 kb) was more than twice as long as haploblocks of any autosome.

**Figure 3.**
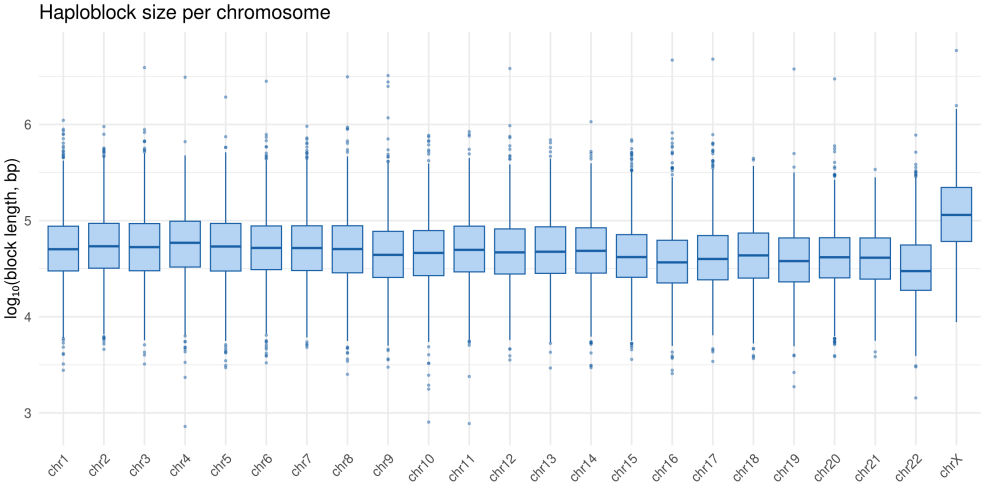
Haploblock length distribution per chromosome. Box plots represent the distributions of haploblock lengths for 22 autosomes (chr1-22) and chromosome X (chrX). Boxes span the interquartile range (IQR; 25-75th percentile) with the median, whiskers extending to 1.5x of the IQR, and outliers beyond the whiskers. The x-axis is ordered by chromosome number, the y-axis shows haploblock length (log_10_-transformed).

Overall, larger chromosomes tended to be divided into more haploblocks. However haploblocks were not correspondingly longer on larger chromosomes and were visibly longer on chromosome X, which recombines only in females. This suggests that the discretization into haploblocks was indeed driven by recombination, rather than chromosome size itself.

### Haploblock sequences capture common and rare haplotypes

We examined the haplotype diversity within haploblocks based on the number of clusters. For each haploblock, similar haplotype sequences were grouped together into the same cluster, therefore the number of clusters reflects how many distinct haplotypes were present in the haploblock. The number of clusters varied widely across chromosomes, ranging from 2019 for chromosome 14 to approximately 5000 for chromosomes 9, 12, 16 and 20 (Supp. Fig. 2).

Part of this heterogeneity was explained by chromosome size, but we also found a positive correlation between haploblock length and the number of clusters genome-wide (Pearson *r* = 0.26) and across all 22 autosomes (Pearson *r* between 0.19 and 0.42). However, a large part of the variation remained unexplained (Supp. Fig. 3). We hypothesized that the remaining variation might be reflected in haplotype diversity among clusters. To assess this, we examined haplotype frequencies within haploblocks using the rank abundance of cluster sizes, where the size of a cluster is the number of haplotype sequences assigned to it and shows how frequently the corresponding haplotype occurs in the haploblock (Figure 4).

**Figure 4.**
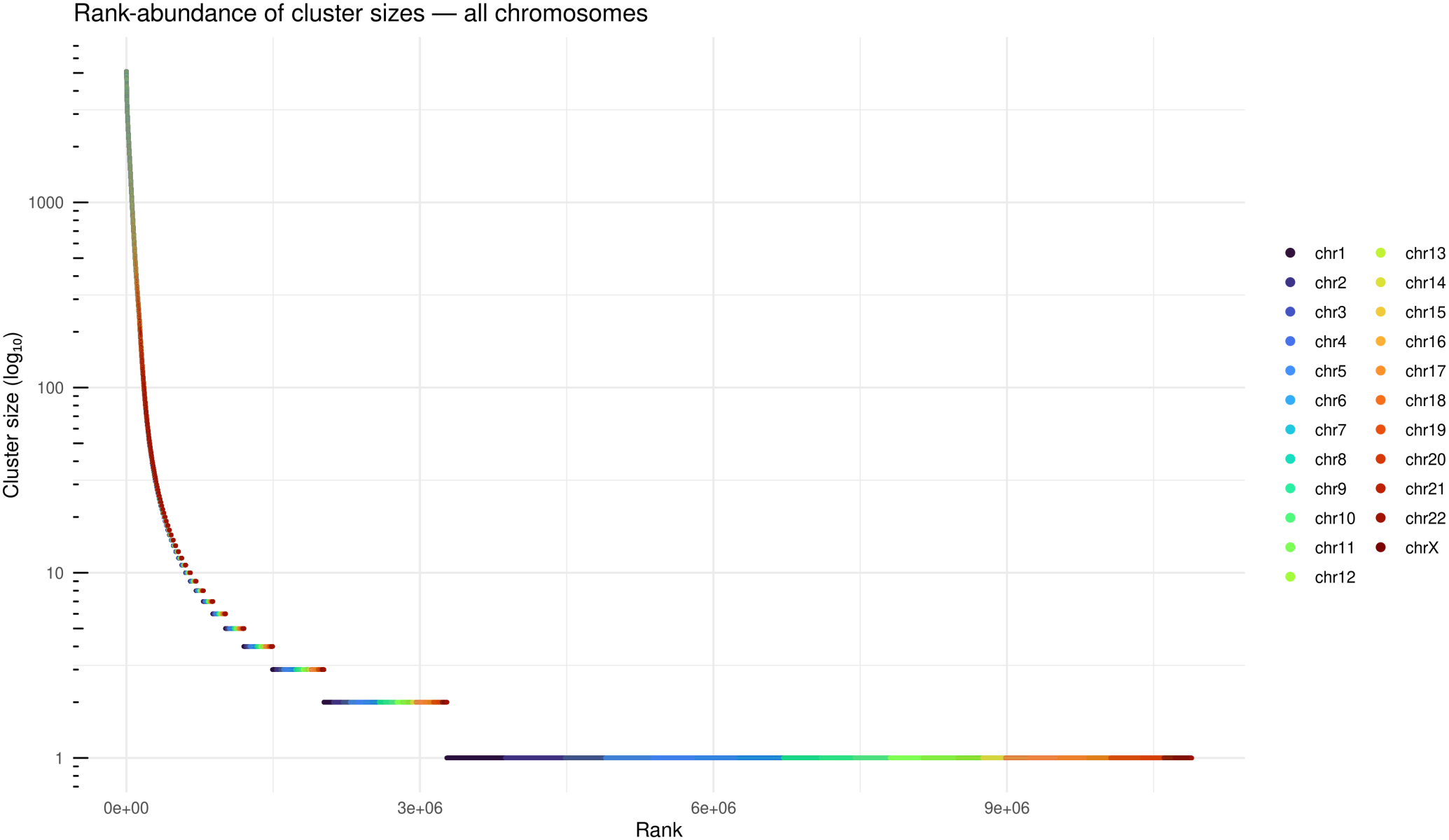
Rank abundance distribution of haploblock cluster sizes. Each point represents a cluster, ranked in descending order of haploblock length (x-axis) and the number of haploblock sequences assigned to that cluster (y-axis, log_10_-transformed). Clusters are ranked in descending size, where the largest clusters have the lowest rank. Clusters, colored by chromosome (chr1-22 and chrX; legend), are shown across all haploblocks genome-wide.

Most haploblock sequences were resolved into a few common clusters: the mean dominant-cluster size was 2502, with most haploblocks (95.5%) containing a dominant cluster with more than 1000 sequences. However, we observed a long tail of small clusters, with the majority of clusters (70.0%) being singletons (cluster size = 1).

To further quantify the haplotype diversity, we calculated the Shannon entropy of cluster sizes (Figure 5, Supp. Fig. 4), which ranges from 0 bits, if all haploblock sequences are assigned to a single cluster, to 12.3 bits, if each sequence is assigned to a distinct cluster. Median entropy varied modestly across autosomal chromosomes and was similarly distributed along each chromosome. Chromosome 13 had the lowest median entropy (2.45), indicating that most individuals carried the same haplotype. In contrast, chromosome 19 had the highest median entropy (3.45), which indicated that the individuals carried more haplotypes. The variation between chromosomes was small relative to the variation within chromosomes: only 1.1% of variance was explained by chromosome identity (*η*^2^ = 0.011, one-way ANOVA).

**Figure 5.**
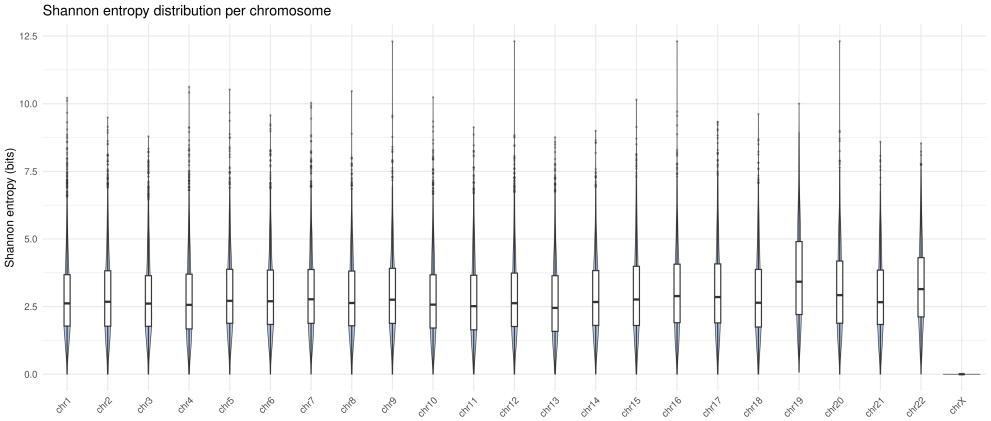
Shannon entropy of cluster sizes per chromosome. Box plots represent the distributions of the Shannon entropy of cluster sizes within haploblocks for 22 autosomal chromosomes (chr1-22) and chromosome X (chrX). Boxes span the interquartile range (IQR; 25-75th percentile) with the median, whiskers extending to 1.5x of the IQR, and outliers beyond the whiskers. The x-axis is ordered by chromosome number, the y-axis shows Shannon entropy.

We also examined the most frequent haplotypes within haploblocks using cluster dominance (Supp. Fig. 5 and 6), defined as the ratio of the largest cluster size to the number of haplotypes in a haploblock. Cluster dominance ranges from approximately 0 if sequences are distributed evenly across many clusters, and equals 1 if one cluster contains every haplotype in the haploblock. The genome-wide median cluster dominance (0.46) indicated that in the most common haploblock the largest cluster included 46% of haplotypes. Only 0.7% of the variance in cluster dominance was explained by chromosome identity (*η*^2^ = 0.007). Therefore no chromosome was dominated by a small number of common haplotypes.

Overall, haploblocks differed in the number of clusters, and haploblock length reflected only a small part of this variation. Most haploblocks were dominated by a small number of common clusters, and the majority of clusters were singletons. This shows that clustering haploblock sequences captured both common and rare haplotypes in the studied population.

## Discussion

The Genomic Cluster Engine discretizes the genome into haploblocks and groups individuals’ phased haploblock sequences into clusters based on sequence similarity [23]. We applied our pipeline to prepare a dataset with haploblock sequences, clusters and genomic hashes for 2,548 individuals from the 1000 Genomes dataset. The standardized binary format of genomic hashes ensures that haploblock-specific patterns of variant co-occurrence are encoded uniquely and in a memory-efficient way. This provides new opportunities to characterize genomic backgrounds that modify variant effects and to define etiological subtypes using haploblock cluster memberships rather than whole-genome sequences.

We examined the haploblock structure and haplotype diversity in the studied populations using the haploblock sequences and clusters. Our results confirm that the discretization of the genome into haploblocks is indeed based on recombination patterns. Furthermore, the clustering of phased sequences within haploblocks can capture both common and rare haplotypes in the population. Finally, the haplotype diversity was consistent across autosomes, in which a typical haploblock was resolved into a few common haplotypes, although in theory many unique sequences are possible.

The haploblock boundaries, sequences, clusters and genomic hashes for the 1000 Genomes dataset are now easily accessible to the research community at https://data.haploblocks.org and on the Registry of Open Data on AWS.

Genomic hashes will enable the stratification of individuals based on their haplotype backgrounds. They will also make it possible to identify the effect of variants in the context of their haplotype backgrounds. When genomic hashes are linked to phenotypic data, the resulting clusters can be associated with phenotypes and traced back to specific combinations of variants that might contribute to specific etiological subtypes. Moreover, because genomic hashes are encoded in a standardized format, they can be produced from other datasets or biobanks using our pipeline and compared directly with other genomic hashes to improve the statistical power of such analyses. Therefore we are working on a solution to add individuals from any biobank or dataset to relevant clusters, without the need to re-run the pipeline. This will allow both private and public datasets to be integrated without the need to share whole-genome sequencing data.

## Data and software availability

The haploblock boundaries, sequences, clusters and genomic hashes for the 1000 Genomes Project are available under the CC0 license at https://data.haploblocks.org and on the Registry of Open Data on AWS at https://registry.opendata.aws (s3 bucket: s3://haploblocks/).

The optimized Genomic Cluster Engine is available under the MIT license on GitHub: https://github.com/collaborativebioinformatics/Genomic_Cluster_Engine.

Configuration templates and dependencies (SAMtools, BCFtools [29], HTSlib [30], MMseqs2) are documented in the GitHub repository as well.

## Author contributions statement

JK: Conceptualization, Formal analysis, Investigation, Methodology, Software, Writing – original draft, Writing – review & editing; MM: Data curation, Formal analysis, Investigation, Software, Visualization, Writing – review & editing; BB: Conceptualization, Funding acquisition, Methodology, Resources, Supervision, Writing – review & editing.

## Acknowledgments

We acknowledge the Danish Centre for AI Innovation (DCAI) for providing access to the AI supercomputer Gefion, and the Danish National Genome Center (DNGC) for computational resources. We also thank NVIDIA, Inc. for providing computational resources.

We thank the AWS Open Data Sponsorship Program for supporting the storage and distribution of the dataset.

MM acknowledges the support of the MDxCORE Department, Rigshospitalet.

We thank Hetvi Jethwani, Krzysztof H. Banecki and Dariusz Plewczynski who contributed to the development of the Genomic Cluster Engine. The initial work on this project was supported by ELIXIR, the research infrastructure for life science data, and conducted at the ELIXIR BioHackathon Europe. We thank the organizers and the European Molecular Biology Laboratory (EMBL) for funding the hackathon.

Clustering was performed using MMseqs2, made publicly available for free by Steinegger and Söding [24].

Figure 1 was created with ChatGPT (https://chatgpt.com) and Claude (https://claude.ai), made publicly available by OpenAI and Anthropic.

