## Supplementary Material for "Generation of a Public Dataset for Etiological Subtyping with the Genomic Cluster Engine"

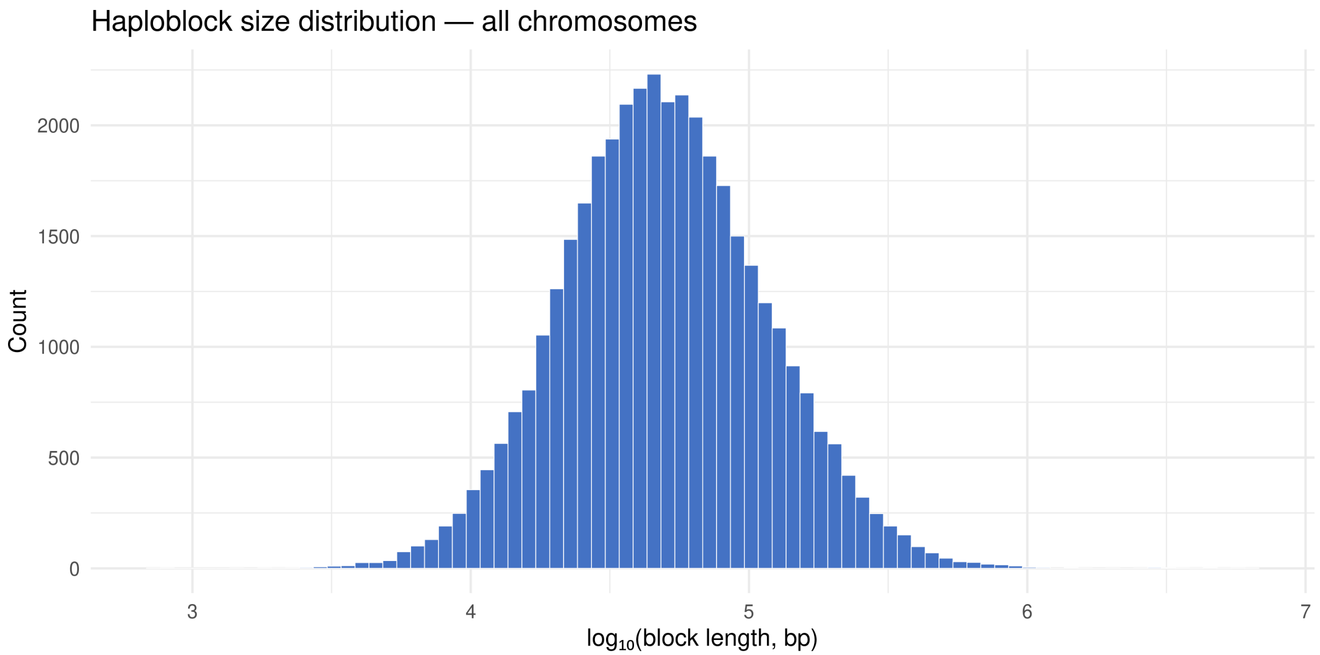


Supp. Fig. 1: **Haploblock length distribution across all chromosomes.** The histogram shows the distribution of 39,141 haploblock lengths across all 23 chromosomes (22 autosomes and chromosome X). The x-axis shows $\log_{10}$-transformed haploblock length in base pairs (bin width = [X] $\log_{10}$ units), the y-axis shows the number of haploblocks per bin.

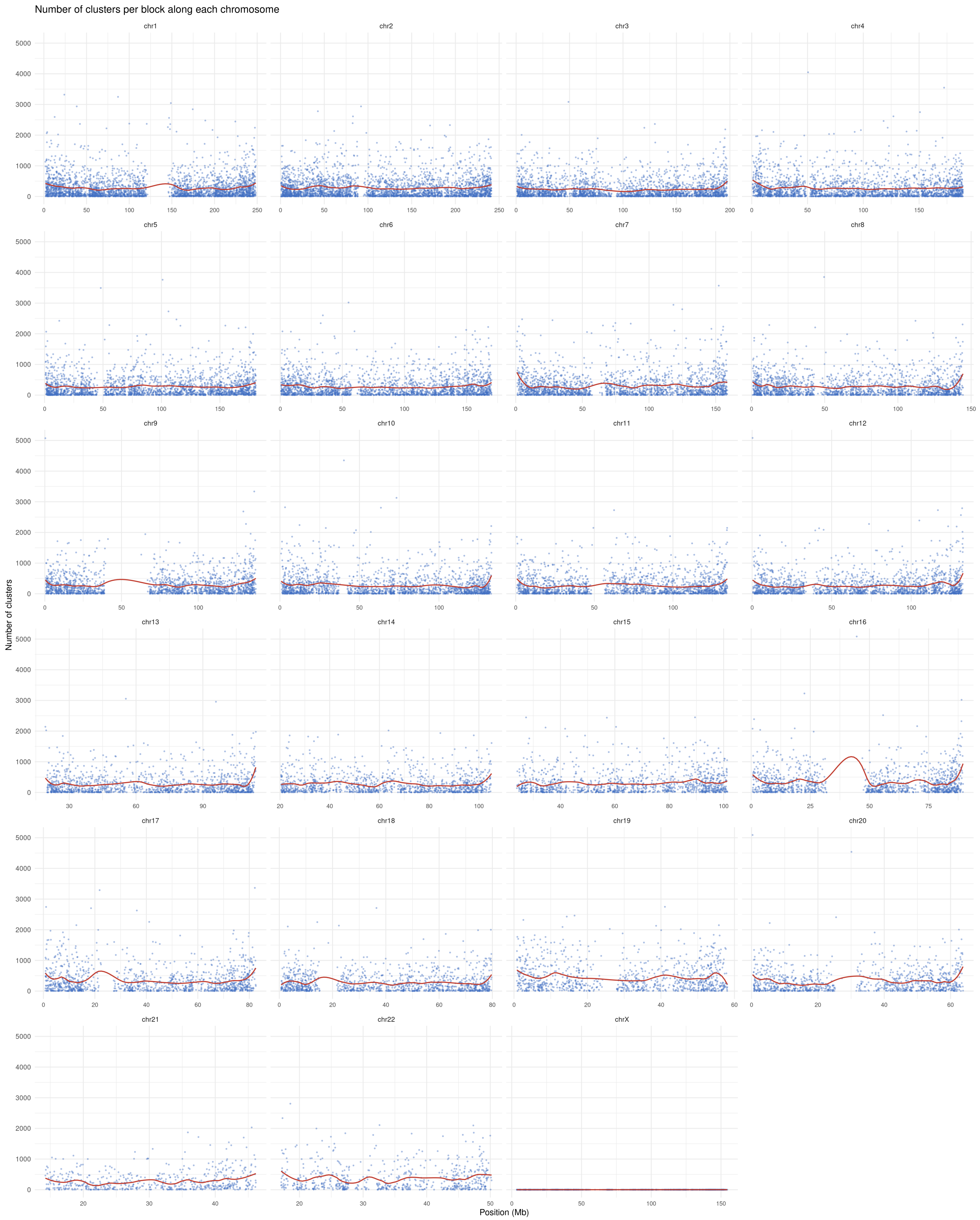


Supp. Fig. 2: **Number of clusters per haploblock along each chromosome.** Each point represents a haploblock, shown for the genomic position on the x-axis and the number of clusters on the y-axis, along each chromosome. The red curves show a smoothed fit. There were no clusters in the centromeres, for which the recombination map and common variants were not available.

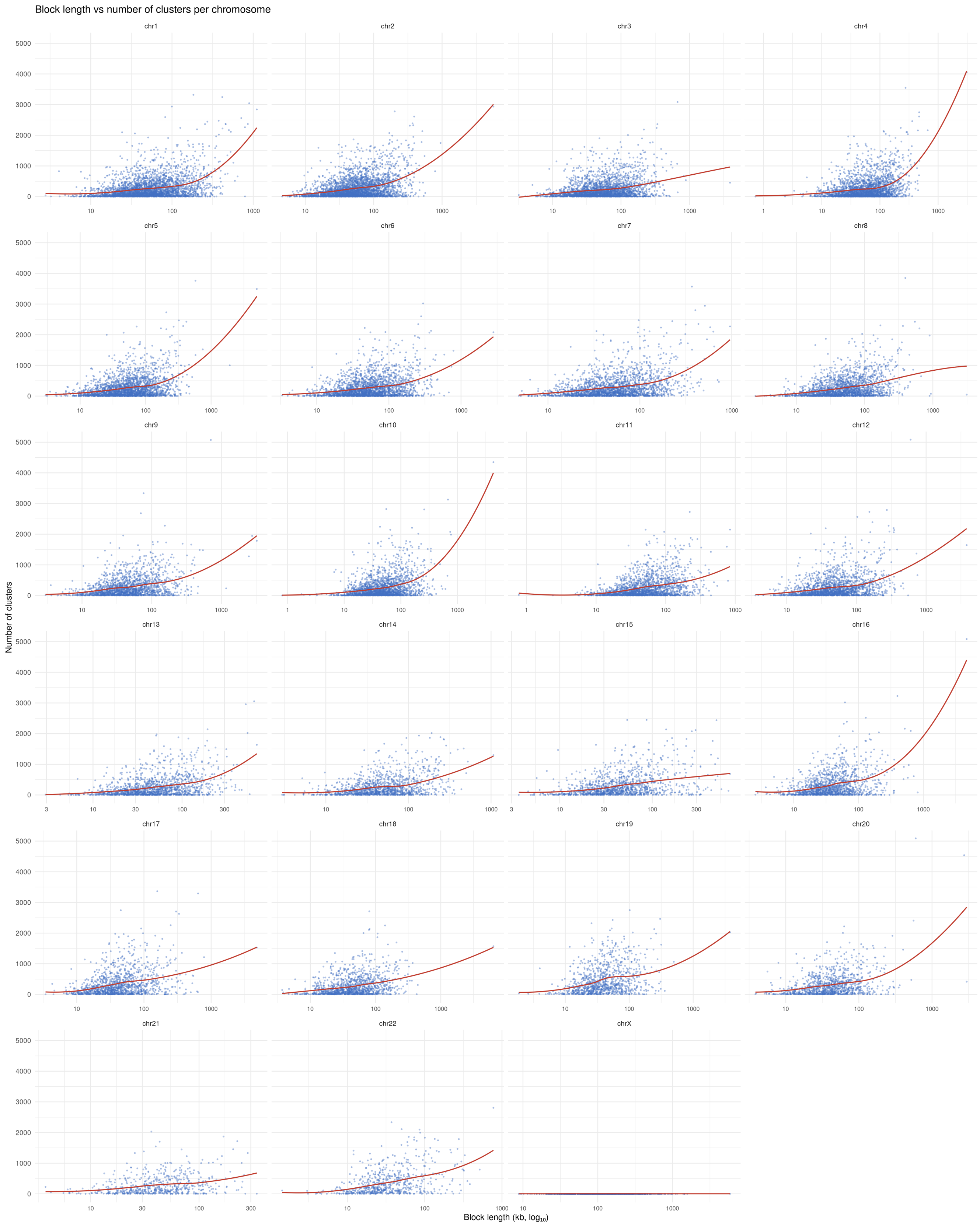


Supp. Fig. 3: **Number of clusters per haploblock as a function of haploblock length along each chromosome.** Each point represents a haploblock, ranked in the increasing order of haploblock length in kb ($\log_{10}$-transformed) on the x-axis and the number of clusters on the y-axis, along each chromosome. The red curves show a smoothed fit.

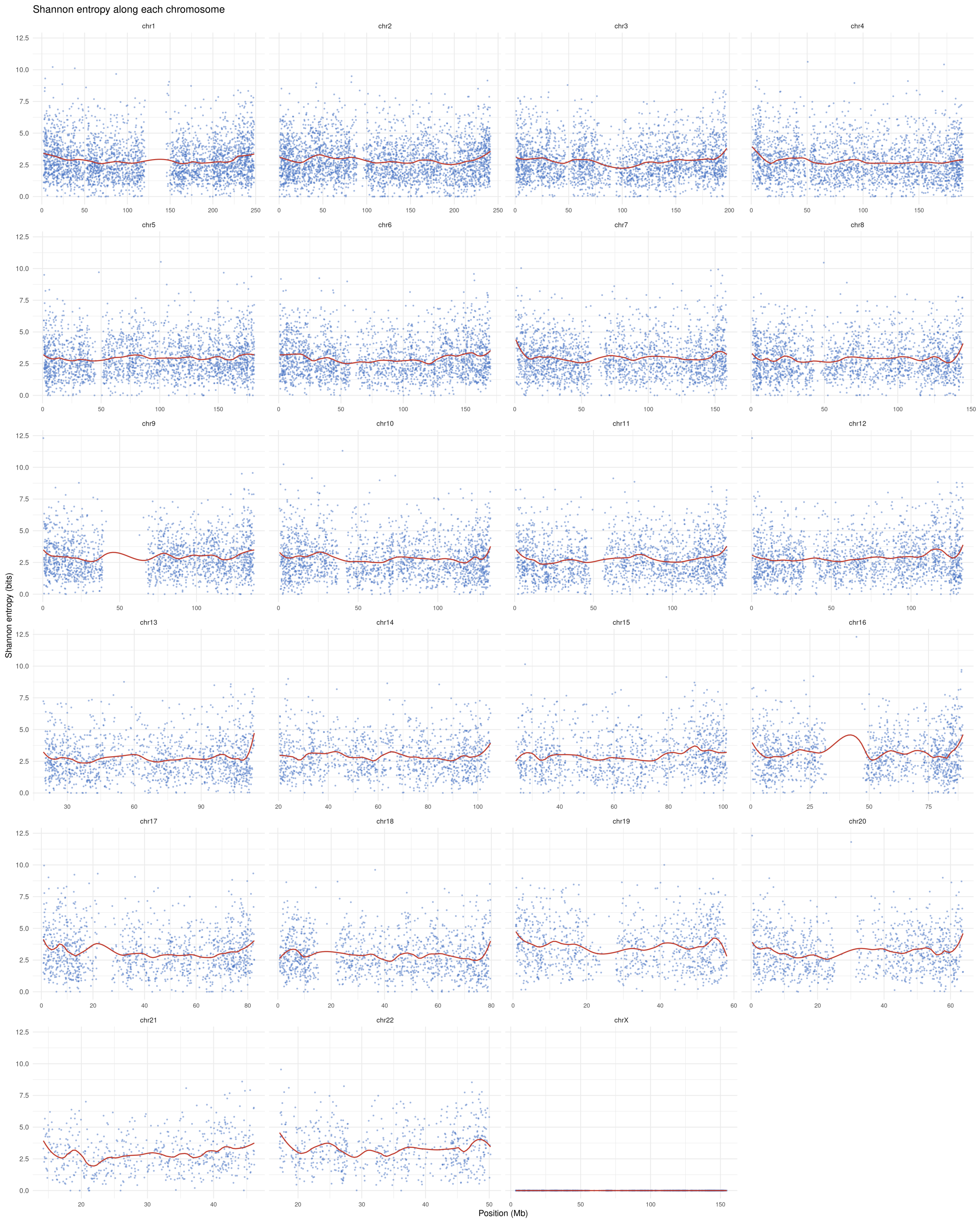


Supp. Fig. 4: **Shannon entropy of cluster sizes along each chromosome.** Each point represents a haploblock, shown for the genomic position on the x-axis and the Shannon entropy of cluster sizes on the y-axis, along each chromosome. The red curves show a smoothed fit. There were no clusters in the centromeres, for which the recombination map and common variants were not available.

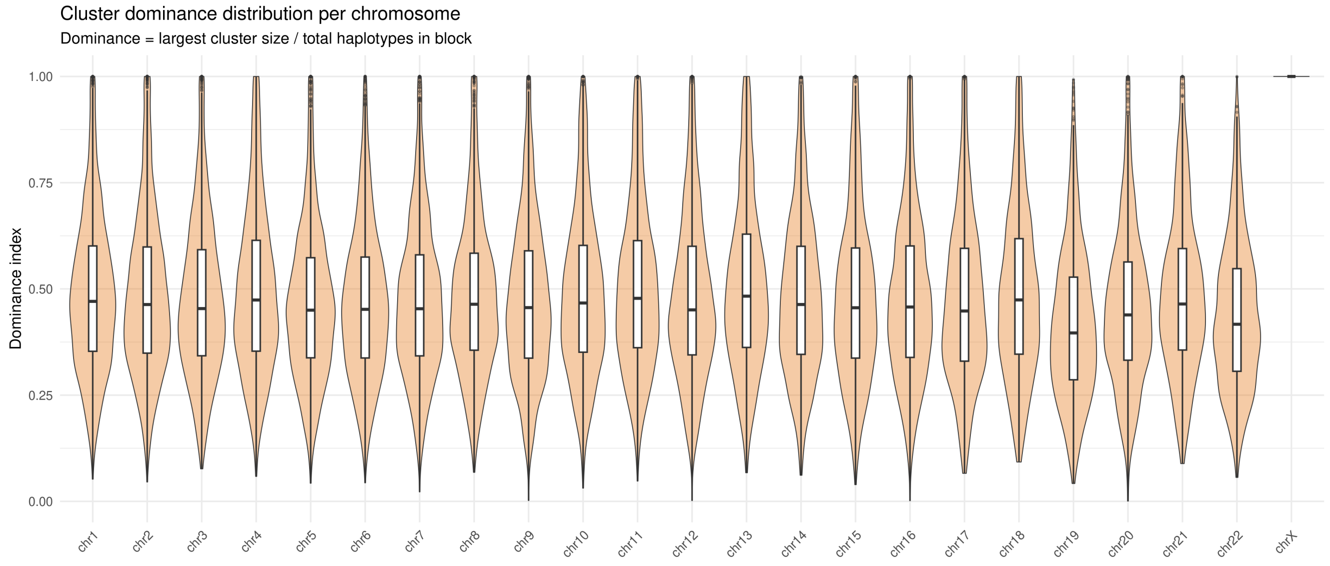


Supp. Fig. 5: **Cluster dominance per chromosome.** The distributions show cluster dominance for 22 autosomal chromosomes (chr1-22) and chromosome X (chrX). Boxes span the interquartile range (IQR; 25-75th percentile) with the median, whiskers extending to 1.5x of the IQR, and outliers beyond the whiskers. The x-axis is ordered by chromosome number, the y-axis shows cluster dominance.

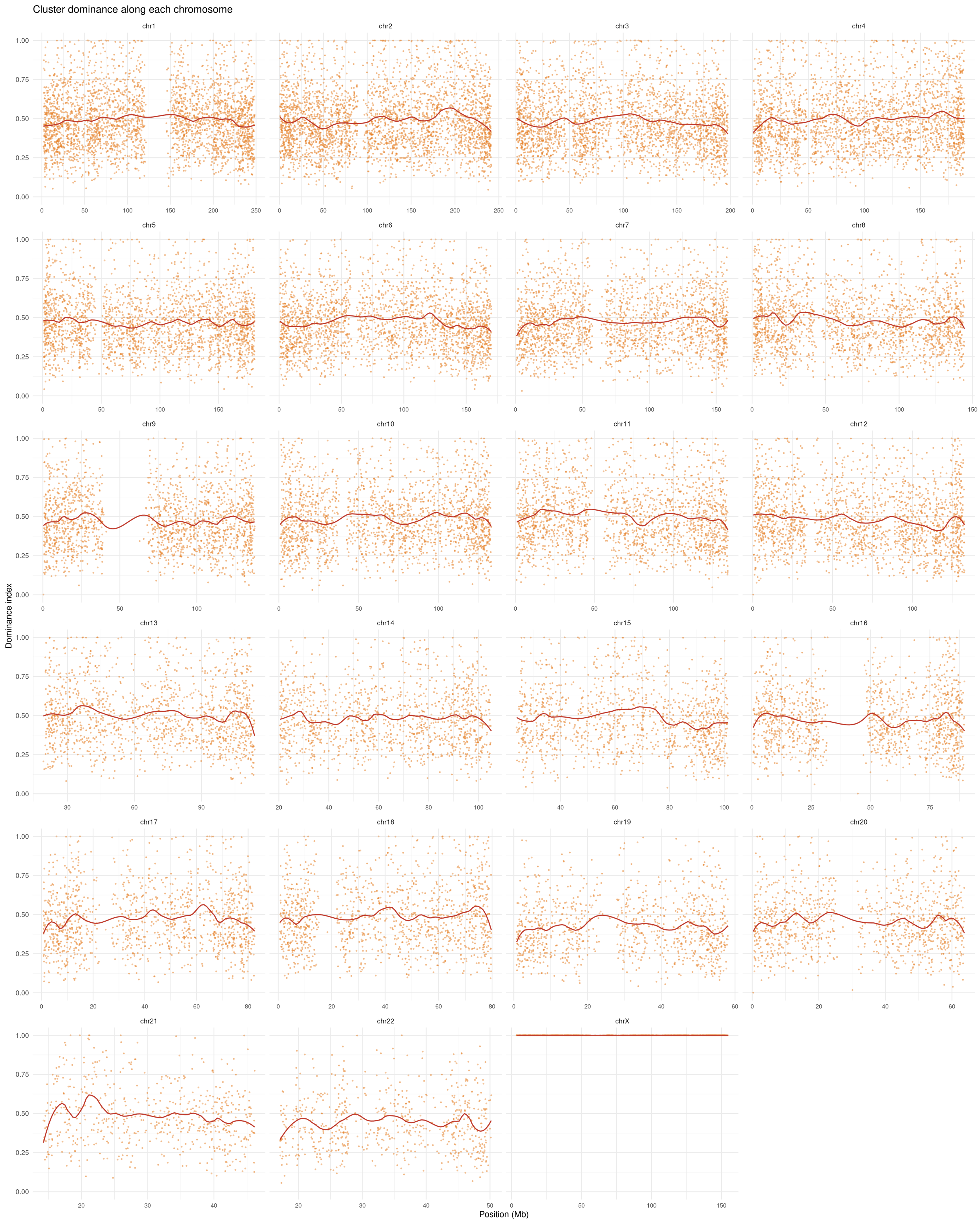


Supp. Fig. 6: **Cluster dominance along each chromosome.** Each point represents a haploblock, shown for the genomic position on the x-axis and the cluster dominance on the y-axis, along each chromosome. The red curves show a smoothed fit. There were no clusters in the centromeres, for which the recombination map and common variants were not available.
